# Comparative analysis of CreER transgenic mice for the study of brain macrophages – a case study

**DOI:** 10.1101/725218

**Authors:** Louise Chappell-Maor, Masha Kolesnikov, Jonathan Grozovski, Jung-Seok Kim, Anat Shemer, Zhana Haimon, Sigalit Boura-Halfon, Takahiro Masuda, Marco Prinz, Steffen Jung

**Author notes:** Corresponding author (S.J.). lead contact (S.J.).

## Abstract

Conditional mutagenesis and fate mapping have contributed considerably to our understanding of physiology and pathology. Specifically, Cre recombinase-based approaches allow the definition of cell type-specific contributions to disease development and inter-cellular communication circuits in respective animals models. Here we compared *Cx*_*3*_*cr1*^*CreER*^ and *Sall1*^*CreER*^ transgenic mice and their use to decipher the brain macrophage compartment as a showcase to discuss recent technological advances. Specifically, we highlight the need to define the accuracy of Cre recombinase expression, as well as strengths and pitfalls of these particular systems that should be taken into consideration when applying these models.

## Introduction

Preclinical studies in laboratory animals have yielded valuable insight into the complexity of multi-cellular organisms, including specific physiological intercellular communication circuits. Moreover, the analysis of dedicated mouse models can provide critical mechanistic insights into specific pathologies. Transcriptome profiling, both at the population and more recently the single-cell level, allows the inference of functional states of specific cell types and definition of responses to stimuli to formulate informed hypotheses. Complementary cell ablation and gene ablation (mutagenesis) strategies enable testing of the latter by probing specific cell and gene functions in physiological *in vivo* contexts. Many of these *in vivo* approaches rely on Cre / loxP-mediated gene manipulation [1], which itself requires cell type-restricted transgenic expression of a constitutive or conditionally active Cre recombinase [2]. Here we provide a case study for the investigation of murine brain macrophages, a particularly challenging area, comparing the performance of two CreER transgenic mouse lines and highlighting pitfalls and strengths of each model.

Brain macrophages have emerged as major players in central nerve system (CNS) physiology and pathophysiology (*Prinz, Jung, Priller, in press*). CNS macrophages comprise parenchymal microglia and border associated macrophages (BAM) that are associated with the meninges, choroid plexus and perivascular niches [3],[4]. Fate mapping studies suggest that with the exception of choroid plexus resident cells, murine CNS macrophages are derived from an early wave of hematopoietic precursors without receiving further contributions from hematopoietic stem cells throughout the life of the organism.

Mice expressing reporter genes or Cre and CreER recombinase transgenes under the control of the promoter driving expression of the CX_3_CR1 chemokine receptor have been widely used to study microglia [5]-[10]. Of note, while *Cx*_*3*_*cr1* is highly expressed in microglia in the CNS, expression is not restricted to these cells, but also found in non-parenchymal macrophages at CNS border locations [3],[11]. Moreover, the *Cx*_*3*_*cr1* promoter is also transiently active during neuroectodermal development [12], and in peripheral immune cells including myeloid precursors [13],[14], as well as DC subsets, T cells and NK cells [15],[16].

Sall1, a member of the Spalt (“Spalt-like” (Sall)) family of evolutionarily conserved transcriptional regulators has emerged as marker that distinguishes *bona fide* parenchymal microglia from BAM [17] and even from parenchymal brain macrophages that seed the brain following engraftment [18] [19]. Accordingly, *Sall1*^*CreER*^ animals [20] were proposed to be more specific for the genetic manipulation of microglial cells [17],[21]. Earlier studies though reported *Sall1* expression also for other glial cells in the CNS [22], as well as the kidney mesenchyme [20].

Here we report a comparative analysis of *Cx*_*3*_*cr1*^*CreER*^ and *Sall1*^*CreER*^ transgenic mice for the study of brain macrophages, using RiboTag-based translatome profiling, as well as histological and flow cytometric analyses of reporter animals. Providing a case study for the recommended accuracy assessment of CreER transgenic animals, we discuss advantages and limitations of these models, as well as potentially useful tamoxifen-independent applications.

## Results and Discussion

### Translatome analysis of CNS cells expressing CreER transgenes under Cx_3_cr1 or Sall1 control

We recently used the RiboTag approach [23] to determine the cell specificity of *Cx*_*3*_*cr1*^*Cre*^ and *Cx*_*3*_*cr1*^*CreER*^ mice [12]. Specifically, Cre-mediated deletion of a *loxP*-flanked (‘floxed’) wild type exon induces the expression of a hemagglutinin (HA) epitope-tagged ribosomal subunit (RPL22), thereby permanently marking defined cells. Moreover, HA-tagged ribosomes can subsequently be immunoprecipitated from crude whole tissue extracts with anti-HA antibody-coupled magnetic beads enabling the pull down of cell-type-specific ribosomal attached mRNA.

To define the CNS cell types targeted by *Cx*_*3*_*cr1*^*CreER*^ and *Sall1*^*CreER*^ transgenic mice in an unbiased way that does not rely on cell retrieval, we used the RiboTag approach on brains of *Cx*_*3*_*cr1*^*CreER*^: - and *Sall1*^*CreER*^ *:Rpl22*^*HA*^ mice, 9 weeks post tamoxifen (TAM) treatment (**Fig. 1A**). Specifically, we performed direct immunoprecipitation (IP) with either anti-HA (‘HA IP’) or IgG isotype antibody as control (‘IgG IP’) and compared transcriptome and translatome of sorted microglia (Ly6C/G^−^ CD11b^+^ CD45^int^) by taking all sorted cells (‘Sort’) or performing IP with anti-HA on sorted microglia (‘Sort / HA IP’), respectively. Principle component analysis (PCA) of the RNA-seq data revealed that as expected, the two mouse strains displayed considerable overlap in IgG control IPs from the crude extracts, and in all the sorted samples. In contrast, however, the HA IP samples of the *Cx*_*3*_*cr1*^*CreER*^:- and *Sall1*^*CreER*^*:Rpl22*^*HA*^ brains showed considerable differences (**Fig. 1B**). To investigate the underlying reason for this discrepancy we performed a hierarchical K-means clustering which highlighted four clusters of genes enriched in the HA IPs compared to the IgG IPs (**Fig. 1C**). This identified 4370 differentially expressed genes (fold change>2, p-adj<0.05). Clusters II and III comprised genes that had previously been noted to be associated with microglia, including *HexB, Tmem119, Cx3cr1* and *Sall1*. Notably, although *Cx3cr1* and *Sall1* are heterozygous in the *Cx*_*3*_*cr1*^*CreER*^:- and *Sall1*^*CreER*^*:Rpl22*^*HA*^ mice, respectively, both mRNAs were significantly enriched in both samples (**Fig. 1C, E**). Reduced enrichment of *Cx*_*3*_*cr1* mRNA in *Sall1*^*CreER*^*:Rpl22*^*HA*^ CreER mice is likely related to the abundant expression of HA-tagged ribosomes in oligodendrocytes and astrocytes in these animals, which saturated the system.

**Figure 1.**
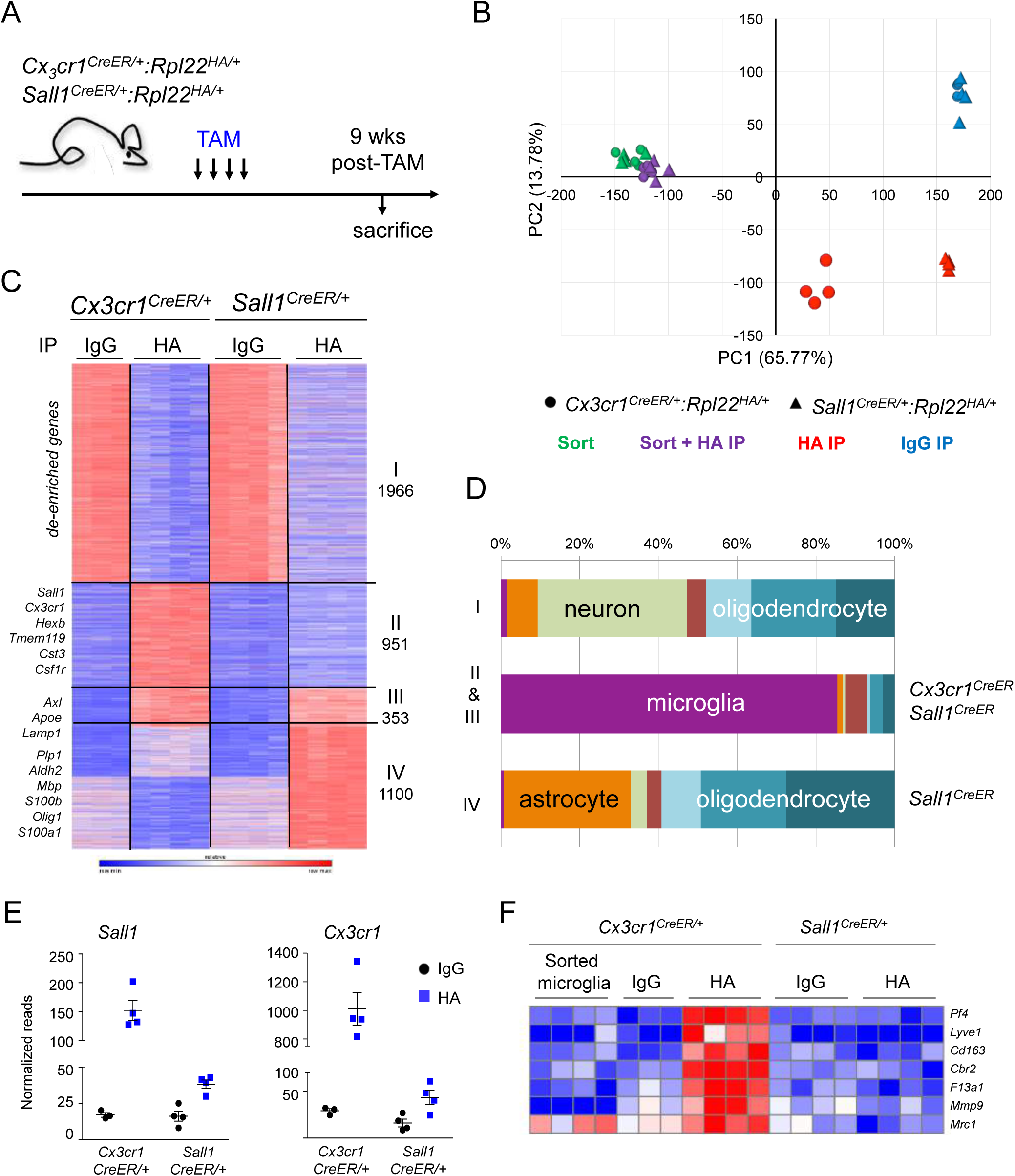
Ribotag analysis reveals alternative specificities of *Cx*_*3*_*cr1*^*CreER*^ and *Sall1*^*CreER*^ driver lines. (A) Scheme of the mouse genotypes used and the treatment timeline (N=4 per genotype). (B) Principle component analysis (PCA) of all samples used in RNA-seq analysis, showing PC1 and PC2. Circles represent *Cx*_*3*_*cr1*^*CreER/+*^*:Rpl22*^*HA/+*^ and triangles represent *Sall1*^*CreER/+*^*:Rpl22*^*HA/+*^, samples immunoprecipitated (IPed) with anti-HA are coloured red (HA IP), IPed with IgG isotype control are coloured blue (IgG IP), directly sorted microglia are coloured green (Sort) and sorted samples which were subsequently IP’ed with anti-HA are coloured purple (Sort + HA IP). (C) Heatmap based on K-means clustering of 4370 genes with a significant difference between at least one group from directly IPed samples only (*i.e.* HA IP and IgG IP for each line), fold change>2, p-adjusted<0.05. (N=4 samples per group from four individual mice per genotype, one outlier removed from the *Cx3cr1*^*CreER*^ IgG IP group). (D) Assignment of the differentially expressed genes from (C) by cluster, to published CNS cell expression signatures, displayed as percentage by cluster. (E) Graphs showing normalized reads of *Sall1* and *Cx*_*3*_*cr1*, each dot represents an individual mouse, lines represent mean and error bars represent SEM. A significant difference is seen for *Cx*_*3*_*cr1* expression between IgG IP and HA IP for both genotypes (Student’s t-test p<0.005 for *Cx*_*3*_*cr1*^*CreER*^ and p<0.05 for *Sall1*^*CreER*^) and for Sall1 expression between IgG IP and HA IP for both genotypes (Student’s t-test p<0.005). (F) Heatmap of representative non-parenchymal brain macrophage genes, showing enrichment in the *Cx*_*3*_*cr1*^*CreER*^ HA IP samples, but not in *Sall1*^*CreER*^ HA IP samples nor in sorted microglia samples (without IP) from *Cx*_*3*_*cr1*^*CreER*^ brains.

Surprisingly, HA IP samples of *Sall1*^*CreER*^ brains displayed a prominent additional signature of 1,100 genes (cluster IV), that was absent from the HA IP samples of *Cx*_*3*_*cr1*^*CreER*^ brains. Comparison of this gene list to published CNS cell expression signatures that were based on sorted cell populations [22], revealed that it is comprised of mRNAs associated with neurons, astrocytes and oligodendrocytes (**Fig. 1D**).

As reported earlier, *Cx*_*3*_*cr1*^*CreER*^ transgenic animals display rearrangements in BAM [6],[12],[24], and BAM signature genes such as *Lyve1* and *Pf4* were hence found enriched in HA IP samples of their brain (**Fig. 1F**). In contrast, and in line with an earlier report [17], HA IP samples of *Sall1*^*CreER*^ brains were de-enriched for the BAM signature genes.

Collectively, these data reveal that rearrangement in *Sall1*^*CreER*^*:Rpl22*^*HA*^ mice while sparing BAM, are not restricted to microglia but also include neurons and other glia. This finding corroborates the earlier notion [12] of the validity of the RiboTag approach to determine the accuracy of Cre and CreER transgenic mouse lines.

### Histological and flow cytometric analysis of CNS cells expressing reporter genes following *Cx*_*3*_*cr1* ^*CreER*^ or *Sall1*^*CreER*^ activation

To complement the translatome analysis with histology and flow cytometric analyses, we analyzed *Cx*_*3*_*cr1*^*CreER*^:- and *Sall1*^*CreER*^*:R26/CAG-tdTomato* reporter animals. In line with published data (Goldmann et al., 2013; Goldmann et al., 2016)[12], reporter gene expression in *Cx*_*3*_*cr1*^*CreER*^*:R26/CAG-tdTomato* mice was restricted to microglial cells and BAM, as identified by Iba1 staining (data not shown) [12]. In contrast, brain sections of *Sall1*^*CreE*R^:*R26/CAG-tdTomato* mice stained with IBA1, and *Sall1*^*CreE*R^:*R26/CAG-tdTomato:Cx*_*3*_*cr1*^*GFP*^ mice displayed abundant red fluorescent in parenchymal non-microglial cells along with tdTomato^+^ microglia, as well as tdTomato^+^ cells with neuronal morphology (**Fig. 2, Suppl Fig. 1**). These results were confirmed by a complementary analysis of brains of *Sall1*^*CreER*^*:R26/CAG-YFP* animals, including a quantification, which revealed that reporter gene expression was restricted to parenchymal microglia, absent from macrophages in perivascular niches, meninges and choroid plexus, but also prominent in Sox9^+^ astrocytes (**Suppl Fig. 2**).

**Figure 2.**
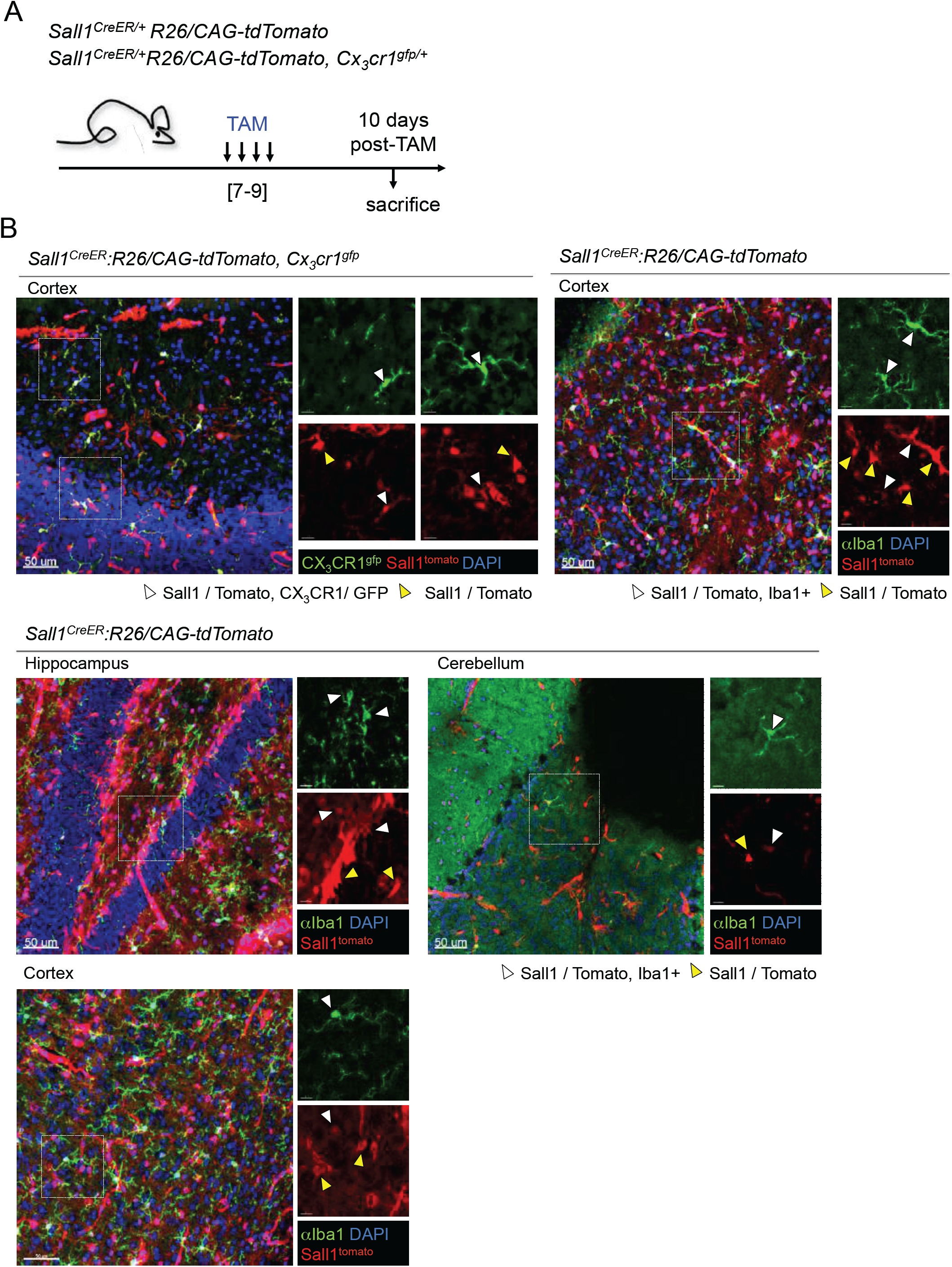
Immuno-histological analyses of *Sall1*^*CreER*^ driver line combined with reporter alleles. (A) Scheme of the mouse genotypes used and the treatment timeline (N=1 per genotype). (B) Representative confocal images of *Sall1*^*CreER*^*:tdTomato, Cx*_*3*_*cr1*^*gfp/+*^ (upper left panel) and *Sall1*^*CreER*^*:tdTomato* mice (upper right and lower panels) of different brain regions (cortex, hippocampus and cerebellum as indicated) stained with anti-Iba1 antibody. *TdTomato*^*+*^ (red) cells co-localizing with GFP^+^/Iba1^+^ microglial cells are indicated by white arrows, and *tdTomato*^*+*^ (red) cells not co-localizing with GFP^+^/Iba1^+^ microglial cells are indicated by yellow arrows. Scale bars represent 50 µm. Representatives of 3 mice per genotype

For the flow cytometric analyses we included the astrocyte-specific *Aldh1I1*^*CreER*^*:R26/CAG-tdTomato* mice as a control [25] (**Fig. 3A**). Macroscopic analysis of brains of *Cx*_*3*_*cr1*^*CreER*^:-, *Aldh1I1*^*CreER*^:- and *Sall1*^*CreER*^*:R26/CAG-tdTomato* mice revealed differential red fluorescent label, with preferential red labelled olfactory bulbs in *Sall1*^*CreER*^*:R26/CAG-tdTomato* mice that was not visible in the two other lines (**Fig. 3B**). Confirming earlier RiboTag results reported for these animals [25], brains of *Aldh1I1*^*CreER*^*:R26/CAG-tdTomato* animals revealed negligible rearrangements in CD11b^+^ CD45^int^ microglia cells (**Fig. 3 C, D, E**). As previously reported [6],[12],[24], *Cx*_*3*_*cr1*^*CreER*^:*R26/CAG-tdTomato* mice showed exclusive reporter gene expression in CD11b^+^ CD45^+^ cells, comprising microglia and BAM. In contrast, *Sall1*^*CreER*^*:R26/CAG-tdTomato* mice displayed a prominent population of labeled CNS cells negative for the pan hematopoietic marker CD45 that likely represent astrocytes and oligodendrocytes (**Fig. 3C, D, E**). Of note, unlike the RiboTag approach or histology, analysis of the CreER transgenic mice crossed to reporter mice by flow cytometry, depends on the isolation of cells and analysis of single cell suspension. Different isolation protocols can yield distinct populations and can hence provide distinct and potentially misleading results.

**Figure 3.**
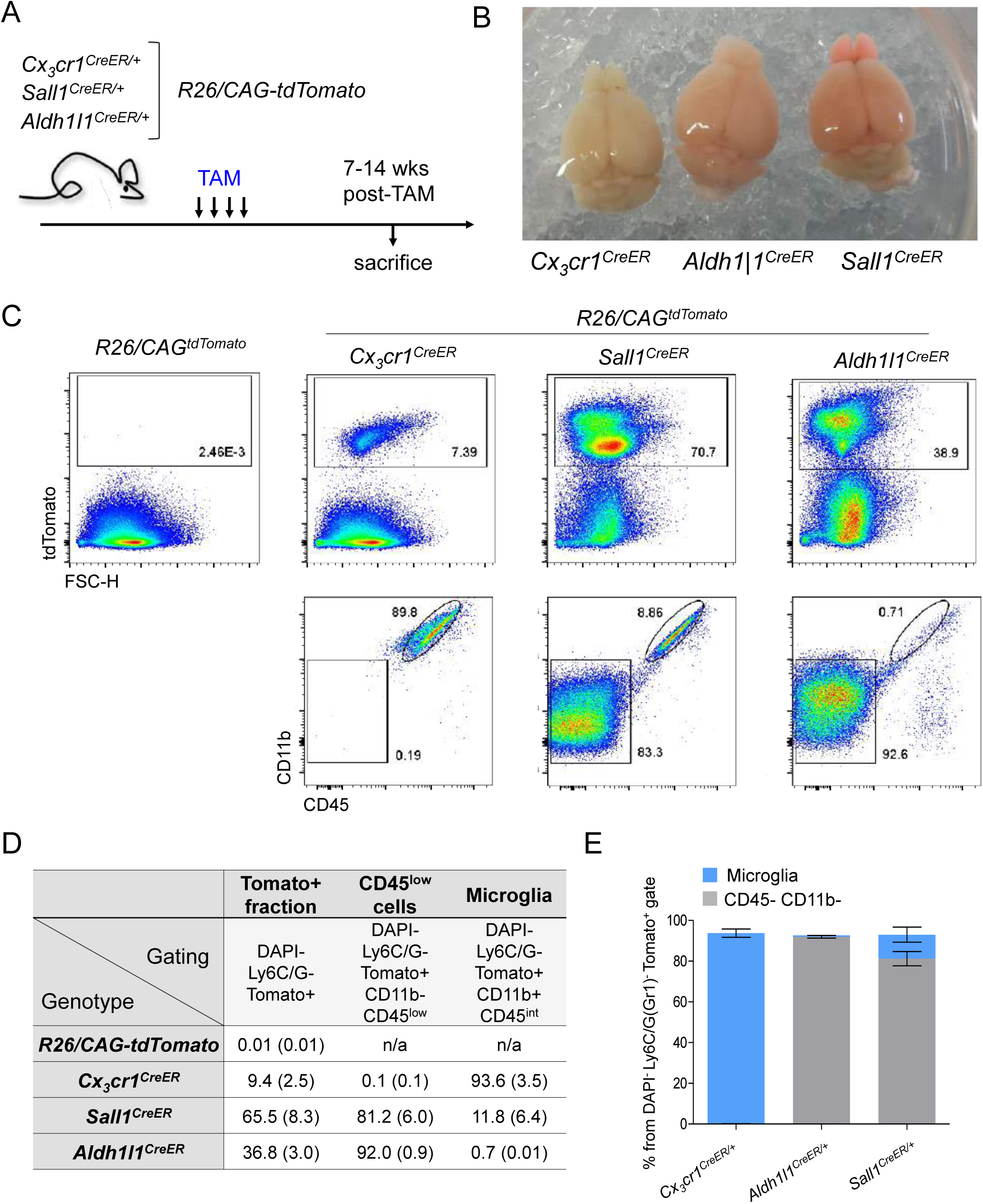
Flow cytometric analyses of *Cx*_*3*_*cr1*^*CreER*^ and *Sall1*^*CreER*^ driver lines combined with reporter alleles. (A) Scheme of the mouse genotypes used and the treatment timeline (N= 3 or 4 per genotype). (B) White light images of *Cx*_*3*_*cr1*^*CreER*^*:-Aldh1|1*^*CreER*^:- and *Sall1*^*CreER*^*:tdTomato* whole brains. (C) FACS plots showing DAPI^−^, Ly6G/C(Gr1)^−^, Tomato^+^ cells (upper panels) separated into microglia (CD11b^+^, CD45^int^) and other glial cells (CD11b^−^, CD45^low^) by flow cytometry (lower panels). (D) Quantification of FACS data from (C), showing mean percentage of parent gate with standard deviations shown in parentheses, for each genotype. (E) Graphical representation of the FACS data from (D) showing the percentage of Tomato^+^ parent gate for microglia and CD45^low^ cells for each genotype. Error bars represent SEM.

### Specific rearrangements in long-lived cells of CreER transgenic animals in absence of tamoxifen treatment

While Cre transgene negative animals never showed activations of the reporter genes, we noted rearrangements in absence of tamoxifen (TAM) treatment when analyzing the various CreER transgenic mice by FACS and immunostainings (**Fig. 4B, C, D**). This is in line with an earlier report on *Cx*_*3*_*cr1*^*CreER*^ mice that harbored a different ‘floxed’ allele [26] and our recent more detailed RiboTag analysis of these animals [12]. In the CreER transgenic approach that was originally introduced by the Chambon group [2], a Cre / estrogen receptor (ER) fusion protein is kept latent in the cytoplasm through binding to heat shock protein 90 (HSP90) [27]. Addition of the estrogen analog TAM frees the recombinase to translocate by virtue of its engineered nuclear translocation signal (NLS) to the nucleus and find the genomic DNA target (**Fig. 4A**). As would be expected, this system is not absolutely tight. Spontaneous nuclear CreER translocation and TAM-independent rearrangements in this model depend on the tightness of the retention and can accordingly be further suppressed by addition of two ER domains [28]. The frequency of TAM independent rearrangements is likely further affected by CreER / HSP90 ratio and the amount of CreER protein present in the given cells, which is governed by the promoter driving the CreER transgene. Another factor, with particular relevance for the long-lived microglia and BAM analyzed in this study, is the time window of CreER expression. TAM independent rearrangements compromise time course experiments and can have direct implications on the interpretation of CreER-based fate mapping studies. Of note however, these spontaneous TAM independent rearrangements are nevertheless specific for the cell types in which the CreER is expressed. Thus, spontaneous rearrangements in *Cx*_*3*_*cr1*^*CreER*^:- and *Sall1*^*CreER*^*:R26/CAG-tdTomato* mice are for instance restricted to the same populations that are also targeted in presence of TAM (**Fig. 4B, C, D, Suppl Fig3B**). If incomplete rearrangements are sufficient for the particular question asked, such as in the case of reporter gene activation for intra-vital imaging, avoidance of TAM treatment, which is known to have considerable side effects [29], could be advantageous.

**Figure 4.**
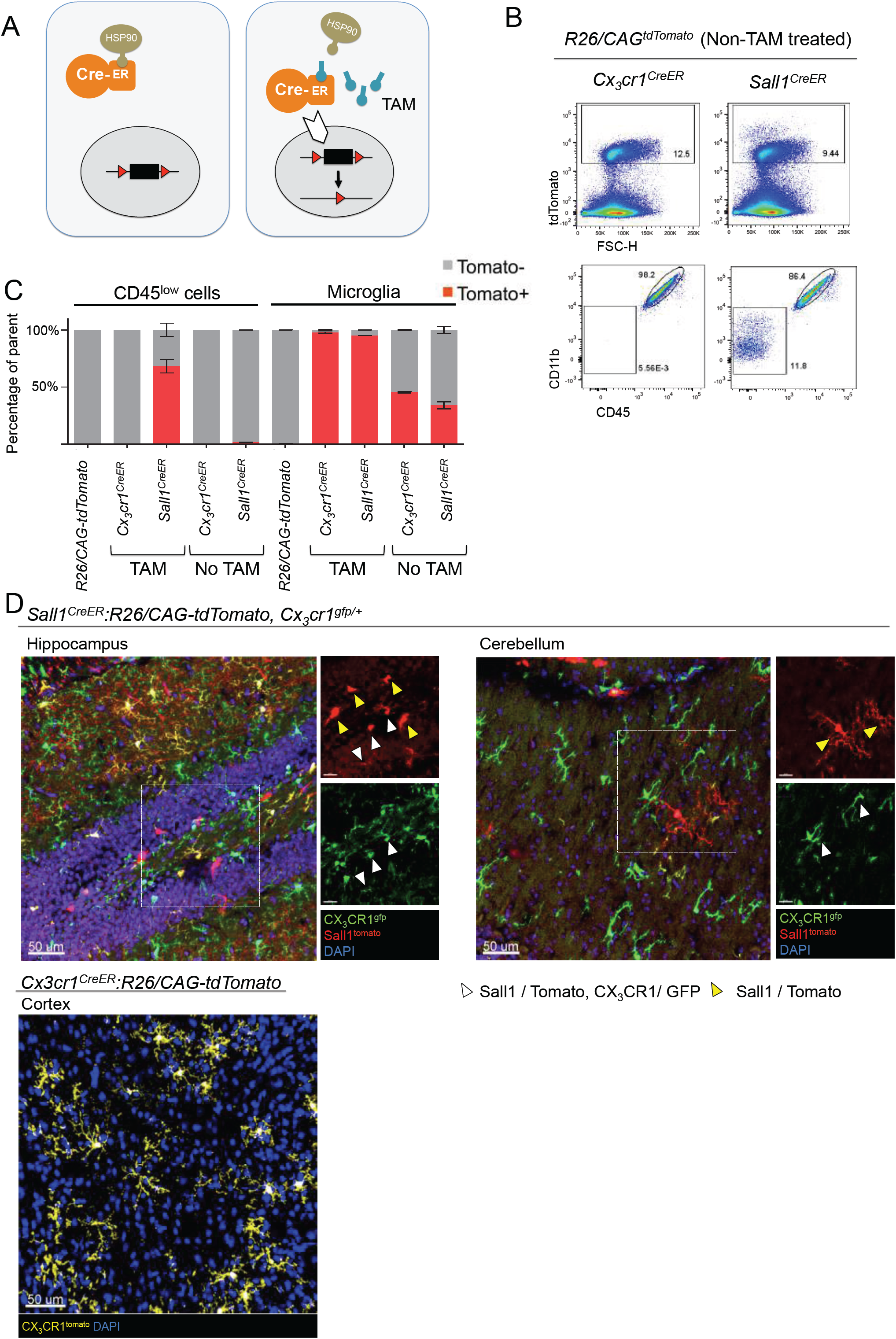
Analyses of the recombinase activity of *Cx*_*3*_*cr1*^*CreER*^ and *Sall1*^*CreER*^ driver lines in the absence of tamoxifen induction. (A) Scheme of mode of action of tamoxifen in inducible CreER driver lines. (B) FACS plots showing DAPI^−^, Ly6G/C(Gr1)^−^, Tomato^+^ cells (upper panels) separated into microglia (CD11b^+^, CD45^int^) and other glial cells (CD11b^−^, CD45^low^) by flow cytometry (lower panels), (N=3 per genotype). (C) Graphical representation of the FACS data from Fig. 3C and 4B combined, showing the percentage of tdTomato^+^ within each cell population (microglia and CD45^low^ cells). Here, TAM treated can be compared to non-TAM treated mice for each genotype. Error bars represent SEM. (D) Representative confocal images of *Sall1*^*CreER*^*:tdTomato:Cx*_*3*_*cr1*^*gfp/+*^ and *Cx*_*3*_*cr1*^*CreER*^*:tdTomato* of different brain regions (hippocampus, cerebellum and cortex as indicated). *TdTomato*^*+*^ (red) cells co-localizing with GFP^+^ microglial cells are indicated by white arrows, and *tdTomato*^*+*^ (red) cells not co-localizing with GFP^+^ microglial cells are indicated by yellow arrows. Scale bars represent 50 µm. (N=1 per genotype).

### Concluding remarks

The present comparative analysis of *Cx*_*3*_*cr1*^*CreER*^ and *Sall1*^*CreER*^ transgenic mice and their application for the study of the murine brain macrophage compartment provides a showcase highlighting strengths and limitations of the available tools. As always, each model has its limitations and particular intricacies the user needs be aware of for the design of the specific experimental set up and the question asked. The study of brain macrophages has been challenging, starting with the observation that the popular *LysM*^*Cre*^ animals, that are often used for the study of peripheral myeloid cells, and were hence used to target microglia, show considerable rearrangements in neurons [30]. Here, we establish that also neither *Cx*_*3*_*cr1*^*CreER*^ nor *Sall1*^*CreER*^ transgenic mice specifically target microglia: *Cx*_*3*_*cr1*^*CreER*^ animals display in addition rearrangements in non-parenchymal CX_3_CR1^+^ CNS macrophages, while *Sall1*^*CreER*^ transgenic mice show prominent recombination in the neuro-ectodermal lineage. These findings are of particular relevance when these animals are used for conditional mutagenesis with ‘floxed’ alleles. Of note though, candidate gens to be targeted can display restricted expression, such as for instance for the hematopoietic lineage, and even phenotypes obtained with the rather promiscuous lines, such as *Sall1*^*CreER*^ transgenic mice, can hence be assignable to cell-type specific deficiencies. Furthermore, imaging studies using CreER mice with the respective reporter animals are less affected by this promiscuity, since they can rely on anatomic or morphologic information for the further definition of cells. Likewise, TAM-independent rearrangements in CreER transgenic mice do not compromise these approaches, since the recombination remains cell-type restricted. TAM-independent rearrangements should however be taken into account when performing fate mapping experiments as they can significantly confound the interpretation of the results.

As for tools targeting specific CNS macrophage subpopulations, it remains unclear whether promoters can be identified that display sufficiently restricted activity, including CNS development and the periphery. Alternatively, transgenic approaches that build on the intersection of two promoters that display overlapping activities could be pursued [31]. This could for example include binary transgenic mice expressing ‘split cre’ fragments under *Sall1* and *Cx*_*3*_*cr1* promoters that according to our data will spare BAM and neuroectoderm, but specifically target microglia. Approaches like these will likely profit from the advent of single cell transcriptomics [11],[32], which can provide valuable data bases for mining (http://mousebrain.org/genesearch.html; http://www.brainimmuneatlas.org). Taken together, our study calls for the informed use of constitutive and conditional Cre recombinase transgenic animal models to avoid technical pitfalls and fully capitalize on their full potential.

## Acknowledgments

The Jung laboratory was supported by the Israeli Science Foundation (887/11), the European Research Council (Adv ERC 340345), and a collaborative network grant of the International Progressive MS Alliance (PMSA). SJ and MP were supported by the Deutsche Forschungsgemeinschaft (DFG) (CRC/TRR167 ‘NeuroMac’).

## Materials and Methods

### Animals

Mice used in this study comprised *Cx*_*3*_*cr1*^*CreER*^ mice (JAX stock # 020940 B6.129P2(C)-*Cx3cr1*^tm2.1(cre/ERT2)Jung^/J); *R26-CAG-tdTomato* mice (B6.Cg-*Gt(ROSA)26Sor*^*tm9(CAG-tdTomato)Hze*^/J) [33], *Sall1*^*CreER*^ mice [20], RiboTag^flox^ mice (JAX stock # 011029 B6N.129-*Rpl22*^*tm1.1Psam*^/J) [23] and Rosa26^YFP^ mice [34]. All animals used in this study were in C57BL/6JOlaHsd background and bred at the Weizmann Institute animal facility. The mice were maintained on a 12 hour light/dark cycle, food and water with provided *ad libitum*. Animals were maintained under specific pathogen-free (SPF) conditions and handled according to protocols approved by the Weizmann Institute Animal Care Committee IACUC as per international guidelines.

### Tamoxifen treatments

To induce gene recombination in CreER transgenic mice, tamoxifen (TAM) was dissolved in warm corn oil (Sigma) and administered orally via gavage for four times every other day. All animals were TAM-treated first at 4–6 weeks of age. Each oral application consisted of 5 mg at a concentration of 0.1 mg/µl. Mice were sacrificed at 7-14 weeks post-TAM treatment as indicated.

### Microglial cell isolation procedure and RiboTag immune-precipitation

Mice were anesthetized with Pental (300 mg/kg, i.p.) and transcardially perfused with PBS. Microglial cells were isolated and sorted as described in [12]. For RNA sequencing of the directly sorted microglia, 10^4^ cells were sorted into L/B buffer (mRNA DIRECT isolation kit, Invitrogen) and frozen at -80°C. For the Sort-IP samples, 3*10^4^ cells were sorted into homogenization buffer (50 mM Tris, pH 7.4, 100 mM KCl, 12 mM MgCl_2_, 1% NP-40, 1 mM DTT, 1:100 protease inhibitor (Sigma), 200 units/ml RNasin (Promega) and 0.1 mg/ml cycloheximide (Sigma) in RNase free DDW) for the Ribotag immunoprecipitation step. Ribotag immunoprecipitation was performed on sorted microglial cells and directly homogenized brain tissue as described in [12]. Ribosome-RNA complexes were eluted from Protein G beads (Invitrogen) into L/B buffer (mRNA DIRECT isolation kit, Invitrogen) and frozen at -80°C.

### RNA-sequencing

Messenger RNA (mRNA) was isolated from the samples using the mRNA DIRECT isolation kit, (Invitrogen). A bulk variation of MARS-seq [35] was used to construct RNA-seq libraries. Final library concentration was measured with a Qubit fluorometer (Invitrogen) and mean fragment size was determined with a 2200 TapeStation instrument. RNA-seq libraries were sequenced using Illumina NextSeq-500. Raw reads were mapped to the genome (NCBI37/mm9) using hisat (version 0.1.6). Only reads with unique mapping were considered for further analysis. Gene expression levels were calculated using the HOMER software package (analyzeRepeats.pl rna mm9 -d <tagDir> -count exons -condenseGenes -strand + -raw) 6. Normalization and differential expression analysis was done using the DESeq2 R-package 30. Differential expressed genes were selected using a 2-fold change cutoff between at least two populations and adjusted pValue for multiple gene testing < 0.05. Gene expression matrix was clustered using k-means algorithm (matlab function kmeans) with correlation as the distance metric. Heatmaps were generated using Genee software.

### Mixed glial cell isolation procedure for FACS analyses

Mice were anesthetized with Pental (300 mg/kg, i.p.) and transcardially perfused with PBS. Mixed glial cells were isolated according to a protocol adapted from [36],[37]. Briefly, brains were dissected, crudely chopped and incubated in a solution containing BSA (Sigma), papain (Worthington) and ovomucoid trypsin inhibitor (Worthington), shaking at 80 rpm for 40 minutes at 37°C. Following the incubation, a solution containing EBSS (Sigma), DNase (Sigma), D(+)-Glucose (Sigma) and NaHCO_3_ was added and homogenates were filtered through a 150µm mesh. Filtrates were subsequently centrifuged at 600g for 5 minutes at 4°C. The cell pellet was resuspended with 30% percoll solution (Sigma) and centrifuged at 600g without acceleration or deceleration, at room temperature for 25 minutes. Next, the cell pellet was washed in PBS and passed through an 80µm mesh, followed by antibody (Ab) labeling and flow cytometric analysis.

### Flow cytometry

Antibodies against CD11b (M1/70), Ly6C/G (Gr-1) (RB6-8C5), CD45 (30-F11) purchased from Biolegend were used at concentration of 1:200 for staining. Analysis was performed on Fortessa (BD Biosciences, BD Diva Software) and analyzed with FlowJo software (Treestar).

### Immunohistology

Mice were anesthetized with Pental (300 mg/kg, i.p.) and transcardially perfused with PBS. Brains were dissected and fixed with 2% PFA overnight at 4°C, and then incubated in 30% sucrose for at least 48h at 4°C. Samples were embedded in optimum cutting temperature compound (OCT), and snap-frozen in isopentane precooled with liquid nitrogen. Sampled were cut into 20 µm sections at -20°C cryostat. Brain sections were washed with PBS, blocked and permeabilized with 1% BSA and 0.3% Triton in PBS at room temperature for 1 h. Brain sections were stained with rabbit anti-Iba1 (019-19741, Wako) overnight at 4°C. Sections were washed three times for 5 min each at room temperature followed by staining with anti-rabbit secondary antibody (711-605-152, Jackson laboratories) for 2 h at room temperature. Nuclei were stained with DAPI (Sigma) for 5 min. Images were acquired with Zeiss LSM 880 confocal microscope and analyzed with Imaris (Bitplane).

## Supplementary Figure legends

**Supplementary Figure 1.**
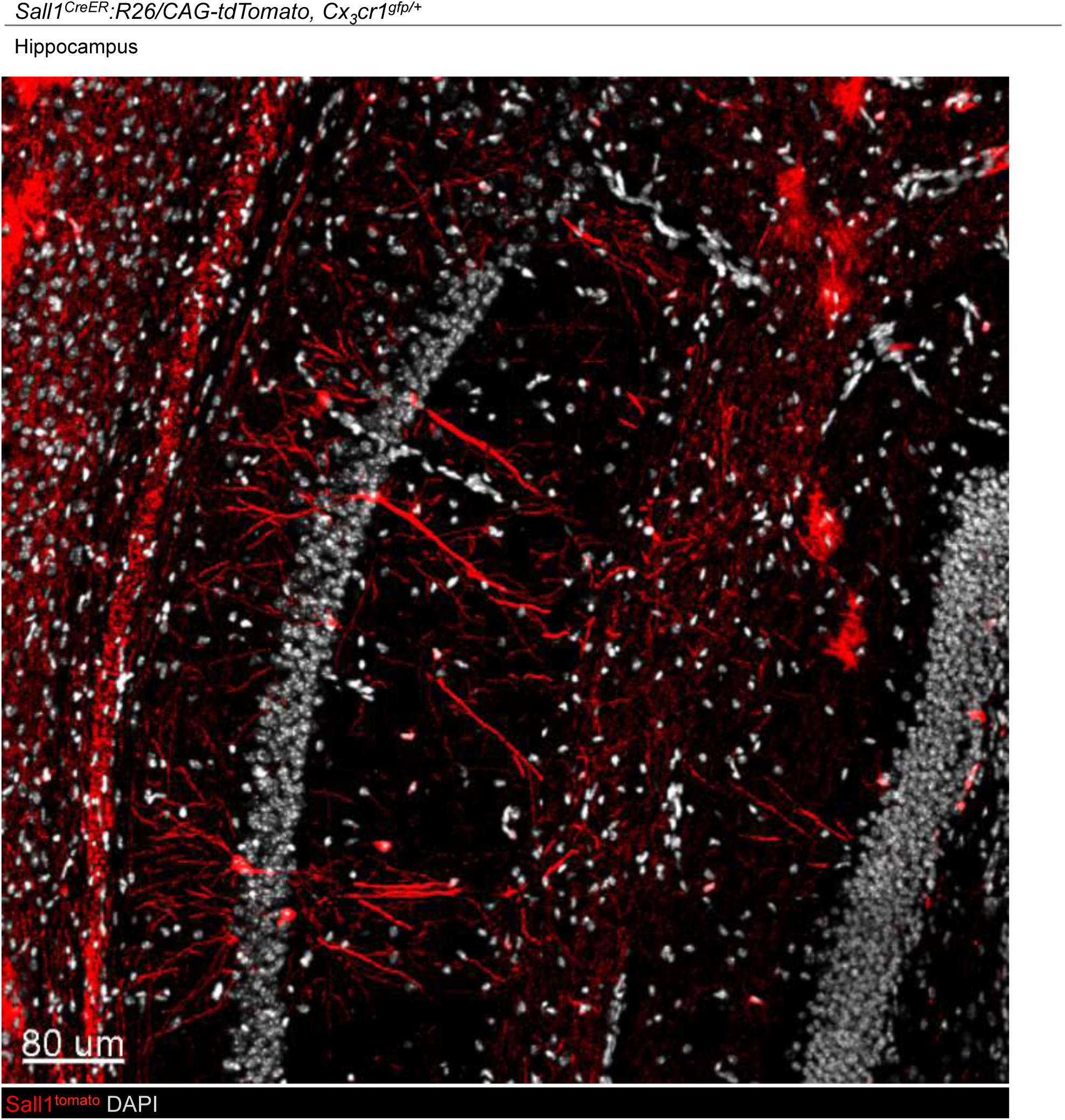
Confocal microscopy analysis of brain hippocampus of *Sall1*^*CreER*^*:R26/CAG-tdTomato* mouse. Red TdTomato reporter gene expressing cells, grey, DAPI

**Supplementary Figure 2.**
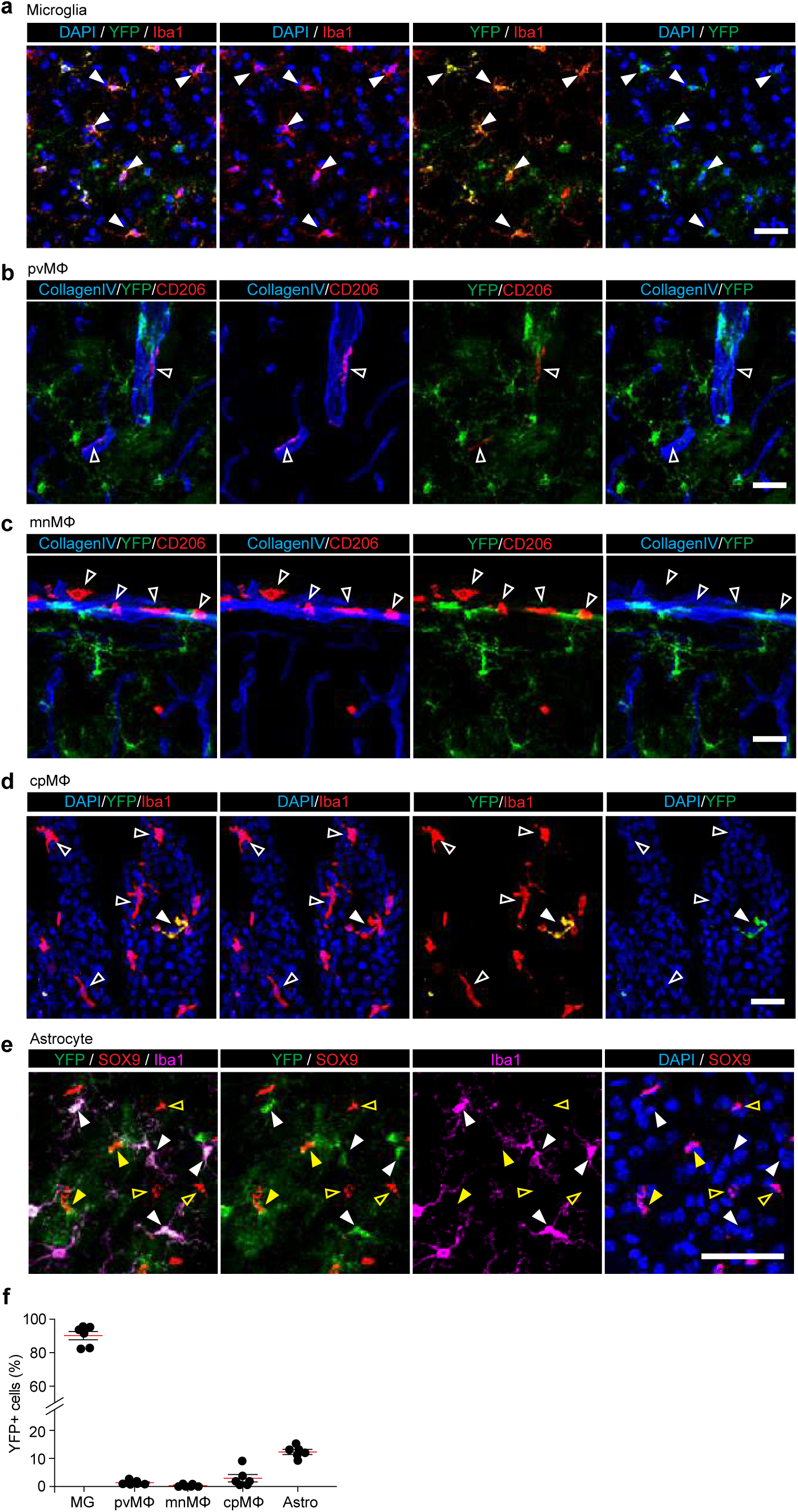
Imunohistochemical analysis of TAM-treated *Sall1*^*CreER*^*:R26*^*YFP*^ *mice*. Representative immunofluorescence images showing YFP labelling in microglia (Iba1, red, **a**), but not in pvMΦ, mMΦ, cpMΦ (CD206 or Iba1, red, **b-d**) and astrocytes (SOX9, red, **e**) in the cortex of adult *Sall1*^*CreER/+*^*R26*^*yfp/yfp*^ mice. Filled or blank white arrowheads in **a**-**d** indicate positive or negative for YFP, respectively. Filled white arrowheads in **e** indicate double positive for Iba1 and YFP, and filled or blank yellow arrowheads show single positive or negative for YFP in SOX9^+^ astrocytes, respectively. Scale bars: 10 µm. Representative images from 6 mice are shown.

**Supplementary Figure 3.**
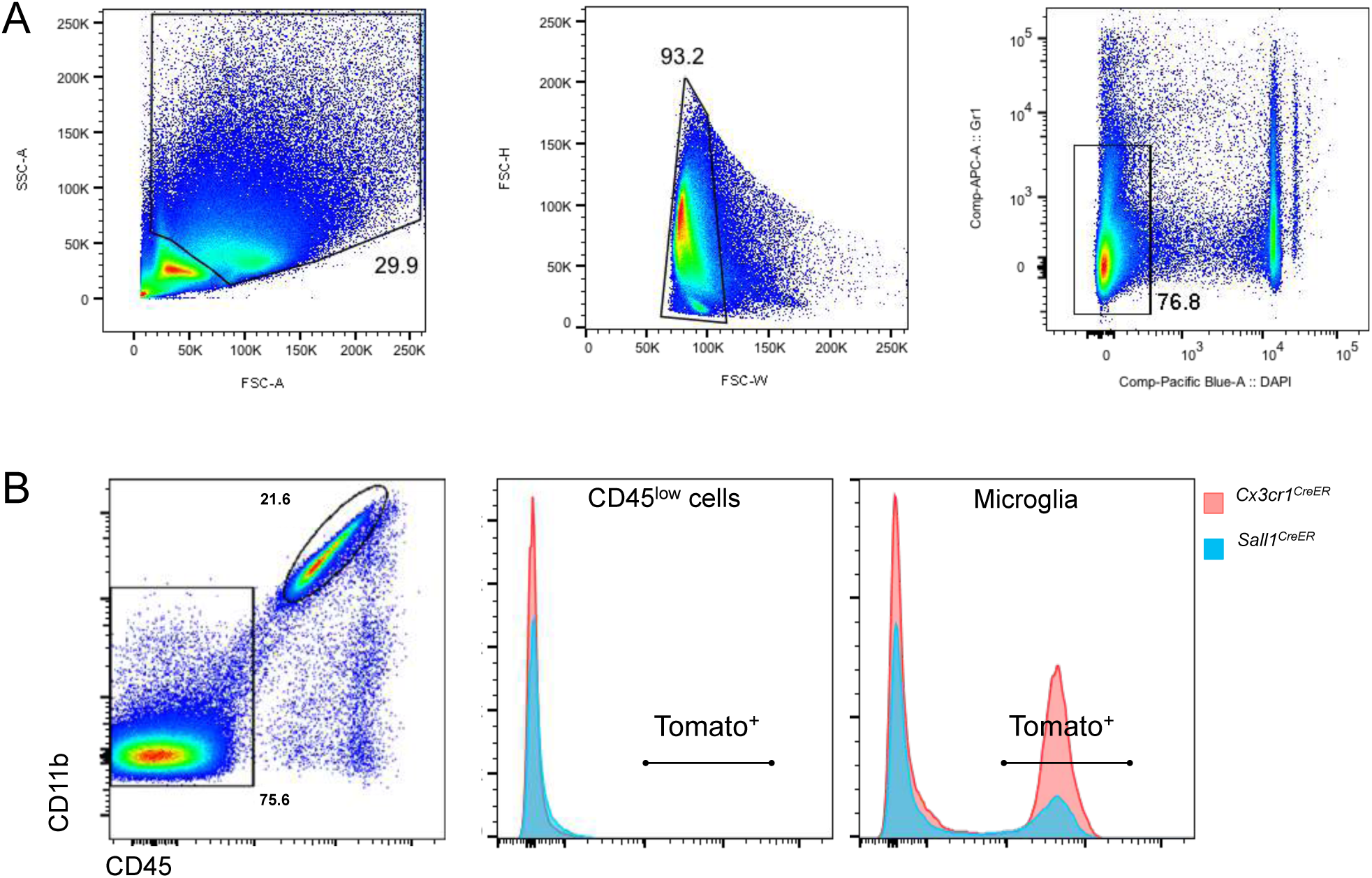
Gating strategy for flow cytometric analysis and quantification. (A) Gating strategy for Figures 3C, 4B and 4C. (B) Gating strategy for quantification of Tomato^+^ cells with CD45^low^ and microglia gates in Figure 4C. Histograms represent tomato labeling in animals not treated with Tamoxifen.

## References

1. Sauer B, Henderson N. Site-specific DNA recombination in mammalian cells by the Cre recombinase of bacteriophage P1. Proc. Natl. Acad. Sci. U.S.A. 1988; 85:5166–5170.DOI: 10.1073/pnas.85.14.5166.

2. Metzger D, Clifford J, Chiba H, Chambon P. Conditional site-specific recombination in mammalian cells using a ligand-dependent chimeric Cre recombinase. Proc. Natl. Acad. Sci. U.S.A. 1995; 92:6991–6995.

3. Goldmann T, Wieghofer P, Jordao MJC, Prutek F, Hagemeyer N, Frenzel K, Amann L, et al. Origin, fate and dynamics of macrophages at central nervous system interfaces. 2016; 17:797–805.DOI: 10.1038/ni.3423.

4. Prinz M, Erny D, Hagemeyer N. Ontogeny and homeostasis of CNS myeloid cells. 2017; 18:385–392.DOI: 10.1038/ni.3703.

5. Jung S, Aliberti J, Graemmel P, Sunshine MJ, Kreutzberg GW, Sher A, Littman DR. Analysis of fractalkine receptor CX(3)CR1 function by targeted deletion and green fluorescent protein reporter gene insertion. Molecular and Cellular Biology. 2000; 20:4106–4114.

6. Goldmann T, Wieghofer P, ller PFMU, Wolf Y, Varol D, Yona S, Brendecke SM, et al. A new type of microglia gene targeting shows TAK1 to be pivotal in CNS autoimmune inflammation. Nature Publishing Group. 2013; 16:1618–1626.DOI: 10.1038/nn.3531.

7. Yona S, Kim K-W, Wolf Y, Mildner A, Varol D, Breker M, Strauss-Ayali D, et al. Fate mapping reveals origins and dynamics of monocytes and tissue macrophages under homeostasis. Immunity. 2013; 38:79–91.DOI: 10.1016/j.immuni.2012.12.001.

8. Davalos D, Grutzendler J, Yang G, Kim JV, Zuo Y, Jung S, Littman DR, et al. ATP mediates rapid microglial response to local brain injury in vivo. Nat Neurosci. 2005; 8:752–758.DOI: 10.1038/nn1472.

9. Nimmerjahn A, Nimmerjahn A, Kirchhoff F, Helmchen F. Resting microglial cells are highly dynamic surveillants of brain parenchyma in vivo. Science. 2005; 308:1314–1318.DOI: 10.1126/science.1110647.

10. Tay TL, Mai D, Dautzenberg J, Fernández-Klett F, Lin G, Sagar Datta M, et al. A new fate mapping system reveals context-dependent random or clonal expansion of microglia. Nat Neurosci. 2017; 20:793–803.DOI: 10.1038/nn.4547.

11. Van Hove H, Martens L, Scheyltjens I, De Vlaminck K, Pombo Antunes AR, De Prijck S, Vandamme N, et al. A single-cell atlas of mouse brain macrophages reveals unique transcriptional identities shaped by ontogeny and tissue environment. Nat Neurosci. 2019; 22:1021–1035.DOI: 10.1038/s41593-019-0393-4.

12. Haimon Z, Volaski A, Orthgiess J, Boura-Halfon S, Varol D, Shemer A, Yona S, et al. Reevaluating microglia expression profiles using RiboTag and cell isolation strategies. 2018; 159:1312.DOI: 10.1038/s41590-018-0110-6.

13. Fogg DK, Sibon C, Miled C, Jung S, Aucouturier P, Littman DR, Cumano A, et al. A clonogenic bone marrow progenitor specific for macrophages and dendritic cells. Science. 2006; 311:83–87.DOI: 10.1126/science.1117729.

14. Liu K, Victora GD, Schwickert TA, Guermonprez P, Meredith MM, Yao K, Chu FF, et al. In vivo analysis of dendritic cell development and homeostasis. Science. 2009; 324:392–397.DOI: 10.1126/science.1170540.

15. Bar-On L, Birnberg T, Lewis KL, Edelson BT, Bruder D, Hildner K, Buer J, et al. CX3CR1+ CD8alpha+ dendritic cells are a steady-state population related to plasmacytoid dendritic cells. Proceedings of the National Academy of Sciences. 2010; 107:14745–14750.DOI: 10.1073/pnas.1001562107.

16. Imai T, Hieshima K, Haskell C, Baba M, Nagira M, Nishimura M, Kakizaki M, et al. Identification and molecular characterization of fractalkine receptor CX3CR1, which mediates both leukocyte migration and adhesion. Cell. 1997; 91:521–530.DOI: 10.1016/s0092-8674(00)80438-9.

17. Buttgereit A, Lelios I, Yu X, Vrohlings M, Krakoski NR, Gautier EL, Nishinakamura R, et al. Sall1 is a transcriptional regulator defining microglia identity and function. 2016; 17:1397–1406.DOI: 10.1038/ni.3585.

18. Shemer A, Grozovski J, Tay TL, Tao J, Volaski A, Süß P, Ardura-Fabregat A, et al. Engrafted parenchymal brain macrophages differ from microglia in transcriptome, chromatin landscape and response to challenge. Nat Comms. 2018:1–16.DOI: 10.1038/s41467-018-07548-5.

19. Bennett FC, Bennett ML, Yaqoob F, Mulinyawe SB, Grant GA, Gephart MH, Plowey ED, et al. A Combination of Ontogeny and CNS Environment Establishes Microglial Identity. Neuron. 2018:1–23.DOI: 10.1016/j.neuron.2018.05.014.

20. Inoue S, Inoue M, Fujimura S, Nishinakamura R. A mouse line expressing Sall1-driven inducible Cre recombinase in the kidney mesenchyme. genesis. 2010; 48:207–212.DOI: 10.1002/dvg.20603.

21. Guttenplan KA, Liddelow SA. Astrocytes and microglia: Models and tools. Journal of Experimental Medicine. 2018; 154:jem.20180200–13.DOI: 10.1084/jem.20180200.

22. Zhang Y, Chen K, Sloan SA, Bennett ML, Scholze AR, O’Keeffe S, Phatnani HP, et al. An RNA-sequencing transcriptome and splicing database of glia, neurons, and vascular cells of the cerebral cortex. J. Neurosci. 2014; 34:11929–11947.DOI: 10.1523/JNEUROSCI.1860-14.2014.

23. Sanz E, Yang L, Su T, Morris DR, McKnight GS, Amieux PS. Cell-type-specific isolation of ribosome-associated mRNA from complex tissues. Proceedings of the National Academy of Sciences. 2009; 106:13939–13944.DOI: 10.1073/pnas.0907143106.

24. Jordao MJC, Sankowski R, Brendecke SM, Sagar, Locatelli G, Tai Y-H, Tay TL, et al. Single-cell profiling identifies myeloid cell subsets with distinct fates during neuroinflammation. Science. 2019; 363:eaat7554–19.DOI: 10.1126/science.aat7554.

25. Srinivasan R, Lu T-Y, Chai H, Xu J, Huang BS, Golshani P, Coppola G, et al. New Transgenic Mouse Lines for Selectively Targeting Astrocytes and Studying Calcium Signals in Astrocyte Processes In Situ and In Vivo. Neuron. 2016; 92:1181–1195.DOI: 10.1016/j.neuron.2016.11.030.

26. Fonseca MI, Chu S-H, Hernandez MX, Fang MJ, Modarresi L, Selvan P, MacGregor GR, et al. Cell-specific deletion of C1qa identifies microglia as the dominant source of C1q in mouse brain. 2017:1–15.DOI: 10.1186/s12974-017-0814-9.

27. Scherrer LC, Picard D, Massa E, Harmon JM, Simons SS, Yamamoto KR, Pratt WB. Evidence that the hormone binding domain of steroid receptors confers hormonal control on chimeric proteins by determining their hormone-regulated binding to heat-shock protein 90. Biochemistry. 1993; 32:5381–5386.DOI: 10.1021/bi00071a013.

28. Verrou C, Zhang Y, Zürn C, Schamel WW, Reth M. Comparison of the tamoxifen regulated chimeric Cre recombinases MerCreMer and CreMer. Biol. Chem. 1999; 380:1435–1438.DOI: 10.1515/BC.1999.184.

29. Hesselbarth N, Pettinelli C, Gericke M, Berger C, Kunath A, Stumvoll M, Blüher M, et al. Tamoxifen affects glucose and lipid metabolism parameters, causes browning of subcutaneous adipose tissue and transient body composition changes in C57BL/6NTac mice. Biochemical and Biophysical Research Communications. 2015; 464:724–729.DOI: 10.1016/j.bbrc.2015.07.015.

30. Orthgiess J, Gericke M, Immig K, Schulz A, Hirrlinger J, Bechmann I, Eilers J. Neurons exhibit Lyz2 promoter activity in vivo: Implications for using LysM-Cre mice in myeloid cell research. Eur. J. Immunol. 2016; 46:1529–1532.DOI: 10.1002/eji.201546108.

31. Hermann M, Stillhard P, Wildner H, Seruggia D, Kapp V, Sánchez-Iranzo H, Mercader N, et al. Binary recombinase systems for high-resolution conditional mutagenesis. Nucleic Acids Research. 2014; 42:3894–3907.DOI: 10.1093/nar/gkt1361.

32. Zeisel A, Hochgerner H, Lönnerberg P, Johnsson A, Memic F, van der Zwan J, Häring M, et al. Molecular Architecture of the Mouse Nervous System. Cell. 2018; 174:999–1014.e22.DOI: 10.1016/j.cell.2018.06.021.

33. Madisen L, Zwingman TA, Sunkin SM, Oh SW, Zariwala HA, Gu H, Ng LL, et al. nn.2467. Nat Neurosci. 2009; 13:133–140.DOI: 10.1038/nn.2467.

34. Srinivas S, Watanabe T, Lin CS, William CM, Tanabe Y, Jessell TM, Costantini F. Cre reporter strains produced by targeted insertion of EYFP and ECFP into the ROSA26 locus. BMC Dev. Biol. 2001; 1:4.DOI: 10.1186/1471-213X-1-4.

35. Jaitin DA, Kenigsberg E, Keren-Shaul H, Elefant N, Paul F, Zaretsky I, Mildner A, et al. Massively parallel single-cell RNA-seq for marker-free decomposition of tissues into cell types. Science. 2014; 343:776–779.DOI: 10.1126/science.1247651.

36. Foo LC. Purification of rat and mouse astrocytes by immunopanning. Cold Spring Harb Protoc. 2013; 2013:421–432.DOI: 10.1101/pdb.prot074211.

37. Rothhammer V, Mascanfroni ID, Bunse L, Takenaka MC, Kenison JE, Mayo L, Chao C-C, et al. Type I interferons and microbial metabolites of tryptophan modulate astrocyte activity and central nervous system inflammation via the aryl hydrocarbon receptor. Nat Med. 2016; 22:586–597.DOI: 10.1038/nm.4106.

